# Single-Cell RNA Sequencing Reveals Endothelial Cell Transcriptome Heterogeneity under Homeostatic Laminar Flow

**DOI:** 10.1101/2020.12.07.414904

**Authors:** Ziqing Liu, Dana L Ruter, Kaitlyn Quigley, Yuchao Jiang, Victoria L Bautch

## Abstract

**Objective:** Endothelial cells that form the innermost layer of all vessels exhibit heterogeneous cell behaviors and responses to pro-angiogenic signals that are critical for vascular sprouting and angiogenesis. Once vessels form, remodeling and blood flow lead to endothelial cell quiescence, and homogeneity in cell behaviors and signaling responses. These changes are important for the function of mature vessels, but whether and at what level endothelial cells regulate overall expression heterogeneity during this transition is poorly understood. Here we profiled endothelial cell transcriptomic heterogeneity, and expression heterogeneity of selected proteins, under homeostatic laminar flow.

**Approach and Results:** Single-cell RNA sequencing and fluorescence microscopy were used to characterize heterogeneity in RNA and protein gene expression levels of human endothelial cells under homeostatic laminar flow compared to non-flow conditions. Analysis of transcriptome variance, Gini coefficient, and coefficient of variation showed that more genes increased RNA heterogeneity under laminar flow relative to genes whose expression became more homogeneous. Analysis of a subset of genes for relative protein expression revealed that most protein profiles showed decreased heterogeneity under flow. In contrast, the magnitude of expression level changes in RNA and protein was coordinated among endothelial cells in flow vs. non-flow conditions.

**Conclusions:** Endothelial cells exposed to homeostatic laminar flow showed increased cohort heterogeneity in RNA expression levels, while cohort expression heterogeneity of selected cognate proteins decreased under laminar flow. These findings suggest that EC homeostasis is imposed at the level of protein translation and/or stability rather than transcriptionally.

## INTRODUCTION

Endothelial cells (EC) drive new blood vessel formation. Once vessels form, EC line all conduits, where they form and maintain a barrier to solute leak, and participate in blood clotting and other homeostatic functions^1-3^. The ability to modulate the relative heterogeneity of an EC cohort in terms of signaling and cell behaviors is critical to vascular function^4^. For example, during sprouting angiogenesis, the formation of new blood vessels from pre-existing vessels, it is important that some EC migrate while others proliferate to expand both the mass and complexity of the network^1^. These differential cellular behaviors result from heterogeneous responses to incoming signals, accompanied by differential gene expression in selected signaling components within an EC cohort^5-7^.

In contrast, EC respond to remodeling cues, including mechanical input from laminar blood flow, to synchronize signaling responses, as they adopt similar cellular morphologies and become quiescent. This change promotes homeostatic functions such as barrier formation, and lesions that perturb EC responses to homeostatic inputs are often lethal and/or lead to diseases such as atherosclerosis^8, 9^. For example, as vessels mature, Notch signaling becomes arterial and relatively equalized in magnitude^10^, while heterogeneous Notch signaling is a hallmark of early network sprouting^11^. EC align along the direction of blood flow and morphologically appear more homogeneous under homeostatic laminar shear stress both *in vivo* and *in vitro*^8^, suggesting a more homogeneous functional status of EC. However, this heterogeneity loss as EC transition from sprouting to homeostasis has not yet been quantified at the genome level.

Genome-wide expression heterogeneity of EC cohorts is not revealed by assays that average signals, such as bulk RNA or protein profiling; thus, we applied single-cell RNA sequencing (RNA-seq) to EC under homeostatic laminar flow to determine transcriptomic heterogeneity, and immunofluorescence with single-cell resolution to interrogate protein expression heterogeneity. Surprisingly, we found overall increased RNA expression heterogeneity under laminar flow, while protein expression heterogeneity decreased, suggesting that flow-mediated EC homeostasis is imposed at the level of protein translation and/or stability rather than transcriptionally.

## MATERIALS AND METHODS

### Cell culture and laminar flow

Human umbilical vein endothelial cells (HUVEC, Lonza, C2519A) were cultured in endothelial growth medium (basal medium-2 (EBM-2, Lonza CC-3156) supplemented with Endothelial Cell Growth Medium (EGM)-2 Bullet Kit (CC-3162; Lonza)). For flow, 5 x 10^5^ passage 4 HUVEC were seeded onto µ channel slides (ibidi, 80176) coated with 5 µg/ml human plasma fibronectin (Roche, 11080938001) in growth medium overnight, then medium was replaced with EBM-2 supplemented with 10% fetal bovine serum (FBS, Gibco, 26140-087). After 4-6 hr, cells were collected for single-cell RNA-seq, fixed for immunofluorescence, or exposed to 15 dyn/cm^2^ laminar shear stress for 72 hr (homeostatic laminar flow, ibidi system). After flow, EC were either collected for single-cell RNA-seq or fixed for immunofluorescence. MEF were isolated as previously described^12^ from E13.5 CD1 mouse embryos, plated into 0.1% gelatin-coated dishes, and cultured in DMEM (Gibco, 11995-065) with 10% FBS. Antibiotics (Gibco, 15240-062) were used in all culture media.

### Single-cell RNA-seq

For each condition, two biological replicates were collected in two independent experiments. HUVEC under flow or non-flow static conditions were digested with 0.05% trypsin-EDTA (Gibco, 25300-054) at 37°C for 3 min, neutralized with 50% newborn calf serum (Gibco, 16010-159) in PBS, collected with a 1 ml syringe, pelleted by centrifugation at 100 *xg* and 4°C for 5 min, washed once with 0.04% bovine serum albumin (BSA) in PBS, filtered with a Flowmi 40µm cell strainer (Sigma, BAH136800040), counted using Countess (Life Technology, AMQAX1000), diluted to 1000 cells/µl with 0.04% BSA in PBS, mixed with 10% MEF in 0.04% BSA/PBS, and loaded onto 10X Genomics Chromium Controller for droplet generation. MEF were added to HUVEC to: 1) determine multiplet rate via mixed-species design; 2) control for technical variability in post-MEF addition steps. HUVEC from 3 ibidi channel slides were combined for each static or flow sample. Viability of HUVEC was >95% via trypan blue staining. 4000 cells (3600 HUVEC, 400 MEF) per sample were loaded. Reverse transcription and library preparation were performed with the Single Cell 3′ Library & Gel Bead Kit v2 according to 10X Genomics’ standard protocol, except that 13 cycles were used for cDNA amplification. Libraries of two flow experiments (Flow1 and Flow2) and two non-flow experiments (Static1 and Static2) were combined and sequenced in one NextSeq500 run using the high output kit at the UNC Translational Genomics Lab (TGL) core. Each sample provided ∼1-1.5 x 10^8^ 150bp paired-end reads. See experimental workflow in **Supp. Fig. IA**.

### Bioinformatic and statistical analysis

Raw reads were mapped to the merged genome of mm10 (mouse) and hg38 (human) using Cell Ranger (10X Genomics). Reads containing the same UMI were collapsed, then assigned to individual cells via barcodes and analyzed in R. The following quality control steps were implemented: 1) mouse-human multiplets containing both mouse and human reads were excluded **(Supp. Fig. IB)**. Total multiplet rate including cross-species multiplets and intra-species multiplets were ∼0.34% based on %MEF and observed cross-species multiplets/sample **(Supp. Fig. IC)**. 2) Outliers with <200 detected genes/cell were excluded as low-quality or debris. 3) Outliers with low (<0.6%) or high (>10%) levels of mitochondria-encoded genes were excluded as cell debris (low) or dying cells (high). 4) Genes whose expression was undetectable in any HUVEC were also excluded, resulting in 20,722 genes in the dataset. A total of 5251 high-quality HUVEC passed all filters, with ∼10,000 UMI counts/cell. Gene expression normalization was performed with the LogNormalize function in Seurat 2.3.4^13^ in R: UMI counts were normalized to 10,000 total counts/cell before ln (normalized counts + 1) was taken. Principle component analysis (PCA, **Supp. Fig. ID-E**) and t-distributed Stochastic Neighbor Embedding (tSNE, **Supp. Fig. IF**) analysis of MEF collected from all 4 samples were performed with Seurat 2.3.4^13^ in R, and cells were well mixed in the low-dimensional representation, suggesting minimal technical batch effect during droplet generation, library preparation, and sequencing.

Differential tests of Gini coefficient (Gini) and coefficient of variation (CV) were performed with DESCEND^14^. To accurately recover true gene expression distribution, genes with overall low expression were filtered out based on: fraction of non-zero counts <5%, number of non-zero counts <20, or mean UMI <0.15. 5370 genes passed the filters, and their true expression distribution in each replicate was deconvolved by DESCEND. Permutation-based differential tests were then performed for Flow1 vs. Static1 and Flow2 vs. Static2, followed by multiple hypothesis testing correction with the Benjamini & Hochberg method in the R package “stats” to obtain FDR-adjusted *p* values. In parallel, differential tests between the flow and the static population as a whole were also performed by including batch as a covariate during distribution recovery. To ensure that the observed increase in RNA expression heterogeneity under flow did not result from subpopulations of transcriptionally flow-unresponsive HUVEC, we also performed differential tests of Gini and CV change between flow-responsive HUVEC and static HUVEC by gating for *KLF2*+, *KLF4*+, or *KLF2*+*KLF4*+ cells in the flow samples (data not shown). The results were highly consistent with results from the ungated flow samples.

Boxplots and violin plots of normalized gene expression were generated by R with the geom_boxplot function of ggplot2 and the VlnPlot function of Seurat, respectively. Differential tests of gene expression levels were performed with the FindMarkers function in Seurat using the Wilcoxon rank-sum test with default settings (log fold-change threshold of 0.25 and Bonferroni corrected p-value threshold of 10^−6^). Curated gene expression level fold change between static and flow conditions was exported from Seurat for bar plots.

Protein expression data are representative of multiple independent experiments. Statistics tests were one-way ANOVA followed by Tukey’s multiple comparison correction and ratio paired t tests in Prism 8.0.1.

### Immunofluorescence

HUVEC in channel slides were fixed in 4% paraformaldehyde in PBS for 10 min, permeabilized in 0.1% Triton X-100 in PBS for 5 min, blocked in 5% donkey serum in PBS for 30 min, sequentially incubated with primary antibodies, secondary antibodies, and DAPI/conjugated PECAM1 antibody (0.04 µg/ml DAPI, Sigma, 10236276001; 1:250 Alexa647-anti-human PECAM1, Biolegend, 303112) in blocking buffer for 1 hr each, then mounted with 80% glycerol in PBS. Cells were washed 3X with PBS for 5 min each between steps, and all incubations were at room temperature. Primary antibodies used include: rabbit anti-HMGN2 (1:500, Cell Signaling, 9437S), mouse anti-NPM1 (1:250, Invitrogen, 32-5200), rabbit anti-DEK (1:500, Abcam, ab221545), rabbit anti-HMGA1 (1:500, Invitrogen, PA5-78007), rabbit anti-ANKRD1 (1:100, Invitrogen, PA5-30548), mouse anti-β-actin (1:1000, Cell Signaling, 3700), goat anti-Grp78 (HSPA5, 1:20, Santa Cruz, sc-1050), guinea pig anti-vimentin (1:100, Progen, GP53), and rabbit anti-SULF1 (1:50, Invitrogen, PA5-63012). Secondary antibody solution contained 1:250 donkey anti-mouse/rabbit/guinea pig Alexa488/555 (Life Technologies). A control sample without primary antibody was included in each experiment. Z stack images were acquired with an Olympus FV3000 confocal microscope (Olympus) at 60X, and total fluorescence intensity (sum of all slices in the Z stack) of each image was quantified using the Fiji version of ImageJ (NIH). For nuclear staining, only nuclear pixels were quantified by creating a mask using the DAPI channel. For cytoplasmic staining, regions of interest (ROI) were manually drawn for each cell based on cell border staining (PECAM1), then pixels within ROIs were quantified. For each independent experiment, >100 cells from 5 images were quantified. Gini coefficient of total fluorescence intensity was calculated with the Gini function in the package ineq in R.

## Data availability

The single-cell RNA-sequencing data that support the findings of this study are available in GEO under the accession number GSE151867.

## RESULTS

### Endothelial cells have increased transcriptomic heterogeneity under laminar flow

EC become more homogeneous when exposed to homeostatic laminar flow, such as is found in the larger arteries of the body, based on the morphological appearance of EC as they align to the shear stress gradient, and the fact that some signaling (i.e., Notch) is less heterogeneous under laminar flow. We hypothesized that flow-stimulated EC morphological changes were accompanied by homogenization of the transcriptome. To examine the relative heterogeneity of EC at the RNA level with homeostatic laminar flow, we applied 15 dyn/cm^2^ laminar shear stress to HUVEC for 72 hr, which leads to strong EC alignment and morphological homogeneity **(Fig. 1A)**. We then performed single-cell RNA-seq on flowed HUVEC and non-flowed HUVEC controls (called “static”) using the 10x Genomics platform **(Supp. Fig. IA)**. Over 20,000 genes were detected in 5251 high-quality HUVEC, with low doublet rate (<0.5%) and good sequencing depth (∼3000 detected genes/cell, **Supp. Fig. IB-F and IIA**, see **Methods** for quality control). Correlation analysis of pseudo-bulk samples assembled from single-cell RNA-seq data showed a much higher correlation between biological replicates than across different conditions, suggesting minimal batch effects and high reproducibility **(Supp. Fig. IIB and IIC)**. Expression analysis revealed significant expression of the EC markers *PECAM1* and *CDH5* (VE-cadherin), and low or undetectable expression of markers of several other lineages, confirming the EC identity of the analyzed cells **(Supp. Fig. IID)**. Upregulation of two known flow-responsive genes, *KLF2* and *KLF4*^15^, was seen in HUVEC exposed to homeostatic laminar flow **(Fig. 1B)**, confirming appropriate transcriptional flow responses under these conditions.

**Figure 1.**
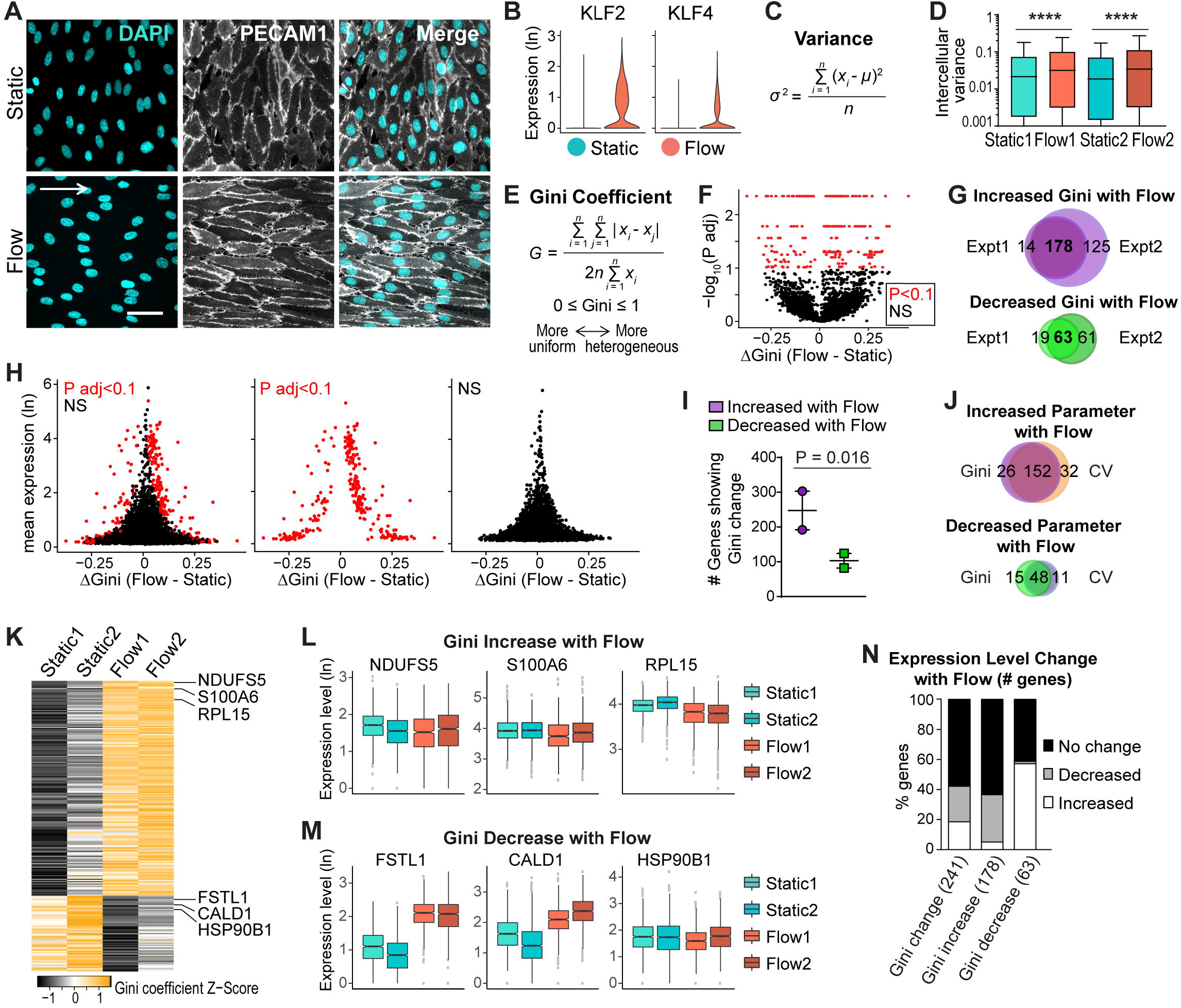
Single-cell RNA-seq shows EC transcriptomic heterogeneity under homeostatic laminar flow. **(A)** HUVEC under indicated conditions labeled for cell-cell borders (PECAM1, white) and nuclei (DAPI, blue). White arrow, direction of flow; Scale bar, 50 µm. **(B-N)** Single-cell RNA-seq of HUVEC under indicated conditions. **(B)** Distribu tion of normalized gene expression of KLF2 and KLF4. **(C)** Variance formula. **(D)** lntercellular variance of HUVEC tran scriptome under indicated conditions. Statistics, one-way ANOVA then Tukey’s multiple comparisons test. ****, p < 0.0001. **(E)** Gini coefficient formula. **(F-J)** Gini was calculated for each gene with DESCEND, and differential test of Gini between conditions was performed. **(F)** Each gene’s Gini change plotted against adjusted p value. Red dots, significant genes (P adj < 0.1); black dots, non-significant genes. **(G)** Venn plots showing gene overlaps of significant Gini change in two independent experiments. **(H)** Each gene’s Gini change plotted against mean expression. (I) Number of genes showing Gini change in each experiment. Statistics, ratio paired t-test. **(J)** Venn plots showing overlap of genes with significant Gini (overlapping genes in G) vs. CV (overlapping genes in Supp. Fig. IIIF) changes. **(K)** Heatmap of Gini coefficient of genes showing significant increased or decreased Gini (overlapping genes in G). **(L-M)** Box plots of overall expression levels of representative genes with Gini increase **(L)** or decrease **(M)**. Hinges correspond to first and third quartiles of expression levels. Gini is indicated by the width of box plots relative to Y-axis. Differential tests of gene expression levels performed with the Wilcoxon rank-sum test (p adj < 10-6). **(N)** Bar plot showing percentage and direc tion of gene expression level changes of genes showing Gini change.

We tested our hypothesis that laminar flow leads to reduced heterogeneity of RNA transcript levels by assessing EC transcriptomic heterogeneity using three different measurements: variance, coefficient of variation (CV), and Gini coefficient (Gini). We first calculated the intercellular variance **(Fig. 1C)** of each gene in each sample.

Surprisingly, a higher overall variance was observed for genes in both replicates of HUVEC under flow compared to static controls **(Fig. 1D)**, suggesting that EC have increased transcriptomic heterogeneity under homeostatic laminar flow. However, since variance can be influenced by gene expression levels, we additionally tested our hypothesis with CV **(Supp. Fig. IIIA)** and Gini **(Fig. 1E)**, both of which are unitless and do not suffer from mean-variance dependence. Since Gini is more robust to extreme outliers than CV^16^, we focused on Gini but also tested CV to determine congruence between the two measurements. We calculated Gini for each gene and performed differential tests with DESCEND^14^, a statistical framework specifically designed for single-cell RNA-seq data **(Fig. 1F)**. We used two strategies to ensure that true biological variation and not batch effect was measured. We first tested each replicate separately and found highly consistent results between replicates, showing largely overlapping lists of genes with significant Gini changes between flow and static conditions **(Fig. 1G)**, suggesting good reproducibility of our data. The second strategy tested the replicates together but with batch as a covariate, and this comparison also led to highly consistent results **(Supp. Fig. IIIB-C)**, suggesting that true equality (Gini) change of RNA expression was measured. We then plotted Gini change between flow and static HUVEC against mean gene expression levels; this comparison suggested that significant Gini changes were not driven by large changes in gene expression, since the number of genes with significant Gini change was not biased towards the top of the y-axis **(Fig. 1H)**. Similar to the variance measurement, a higher number of genes have more heterogeneous expression under flow (178 genes increased Gini) than genes that have more uniform expression under flow (63 genes decreased Gini, **Fig. 1G, I)**, also suggesting increased EC transcriptomic heterogeneity under homeostatic laminar flow. Finally, we calculated CV and performed a differential test as for Gini **(Supp. Fig. IIID-I)**. The results showed that more genes exhibited increased transcriptomic heterogeneity under flow (184 genes increased CV) than decreased transcriptomic heterogeneity (59 genes decreased CV, **Supp. Fig. IIIF**). Moreover, the lists of genes showing significant CV changes between flow and static conditions are largely overlapping with those showing Gini changes **(Fig. 1J, Supp. Fig. IIII)**. In summary, results from three heterogeneity measurements consistently indicate an overall increased transcriptomic heterogeneity of EC under homeostatic laminar flow. These findings refute our hypothesis that loss of heterogeneity at the morphological level is accompanied by homogenization of the RNA transcriptome.

To further examine EC transcriptional heterogeneity in response to laminar flow, we ranked individual genes showing significant Gini changes under flow **(Fig. 1K**). Several genes showing increased Gini under flow by rank were individually plotted by experiment and condition to confirm Gini increase in flow experiments, as evidenced by increased spread along the Y-axis **(Fig. 1L)**. Likewise, several genes ranked for decreased Gini showed decreased spread along the Y-axis in EC exposed to laminar flow **(Fig. 1M)**. We next calculated the percentage of genes whose Gini change are accompanied by expression level changes (**Fig. 1N**). This analysis showed that a majority (58%) of the genes showing significant Gini changes had no overall expression changes. When we further breakdown by the direction of Gini change, genes showing Gini increase with flow were less correlated with overall expression changes than genes showing Gini decrease, with 63% percent of Gini increase genes showing no significant expression change. Overall, this analysis indicates that the increased flow-induced transcriptomic variance in EC resulted from changes unlinked to overall expression level changes.

### EC protein expression becomes more homogeneous under flow

To investigate EC protein expression heterogeneity changes with homeostatic laminar flow, we performed antibody staining for selected EC proteins at single-cell resolution. We tested nuclear-localized proteins to provide more accurate quantification on a per-cell basis, and further selection was based on significant heterogeneity (Gini) change at the RNA level with flow, and antibody availability. Representative images showed more uniform fluorescence intensity per cell under flow compared to static controls **(Fig. 2A-B)**, suggesting decreased protein expression heterogeneity under flow. Quantification of total fluorescence intensity/nucleus, followed by a Gini coefficient calculation, showed decreased protein expression heterogeneity under flow for 4/5 genes, while 1/5 genes showed unchanged heterogeneity under flow **(Fig. 2C)**. These results suggest overall decreased protein expression heterogeneity (Gini) of EC under flow, and they also indicate a decoupling of EC Gini between the RNA transcript and protein level.

**Figure 2.**
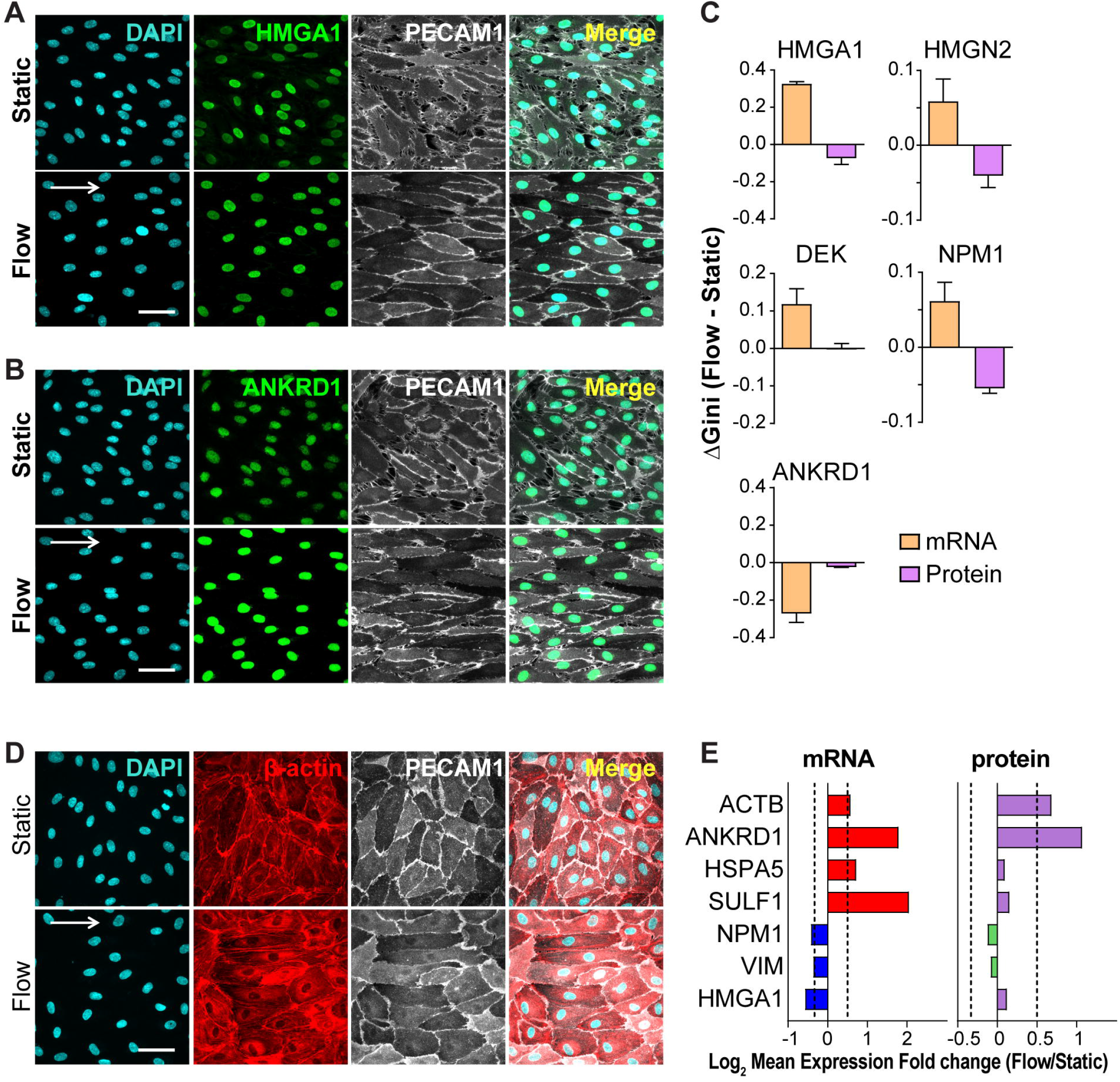
EC protein expression heterogeneity does not follow EC transcriptomic heterogeneity under homeostatic laminar flow. HUVEC immunostained for genes showing significant Gini change **(A-C)** or expression level change **(D-E)** under flow vs. static in single-cell RNA-seq data. Representative images of HMGA1 **(A)**, ANKRD1 **(B)**, and β-actin **(D)** staining, with DAPI (blue, nucleus) and PECAM1 (grey, cell borders). White arrow, direction of flow; scale bar, 50 µm. **(C)** Gini change for tested nuclear genes at the RNA (single-cell RNA-seq) and protein level (immunostaining). Error bars, mean± SEM from multiple replicates. **(E)** Mean expression level fold change for tested genes at the RNA (single-cell RNA-seq) and protein level (fluorescence intensity/cell, representative data of 3 independent experiments). Dotted lines, Fold change = 1.5.

To determine the relationship of overall transcriptomic and protein level changes under flow, we used immunofluorescence to provide semi-quantitative data for proteins whose RNAs showed significant gene expression level changes between flow and static (fold change >1.5) in the single-cell RNA-seq data **(Fig. 2D-E)**. Representative images of β-actin **(Fig. 2D)** and ANKRD1 **(Fig. 2B)** staining showed increased fluorescent intensities under flow compared to static, which are consistent with the single-cell RNA-seq results. Quantification of fluorescence intensity/cell **(right panel, Fig. 2E)** showed that the direction of protein expression level changes between flow and static is consistent with the direction of the RNA expression level changes **(left panel, Fig. 2E)** in 6/7 tested RNA/protein pairs. These results suggest overall agreement of gene expression level changes at the protein and the RNA transcript level in EC exposed to homeostatic laminar flow.

## DISCUSSION

The responses of endothelial cells to laminar flow are crucial to establishing and maintaining vascular homeostasis and quiescence, yet whether the homogenization of morphology and signaling is reflected in global homogenization of transcription profiles was not known. Here we use genome-wide single-cell RNA-seq analysis to show that RNA transcripts become more heterogeneous in expression among a cohort of EC undergoing morphological homogenization in response to flow. In contrast, a sampling of protein expression levels indicates that protein levels become less heterogeneous under homeostatic laminar flow, suggesting that regulation of expression heterogeneity in endothelial cells under flow occurs post-transcriptionally.

Differential gene expression is well-documented between different cell types, but intercellular gene expression heterogeneity within a cell type is relatively understudied due to technical challenges. The recent development of single-cell OMICS and other single-cell techniques have enabled interrogation of global regulation of intercellular heterogeneity, although most studies query expression level changes. Among the few studies focusing on gene expression variation, one provided evidence that regulated heterogeneity but not gene expression levels correlated with epithelial cell differentiation to fiber cells during ocular lens development^17^. DNA methylation heterogeneity across cells was linked to splicing variability during differentiation of induced pluripotent stem cells in another study^18^. Multiplexed RNA *in situ* hybridization revealed high spatial heterogeneity of biomarker gene expression in tumor tissues, suggesting the potential application of spatial heterogeneity as a complementary approach for breast cancer subtype differentiation^19^. Thus intercellular/ interregional gene expression heterogeneity has functional consequences and clinical applications, underscoring its importance.

Endothelial cells are highly specialized in structure and function and remarkably heterogeneous in different tissues and organs^4^. EC transcriptomes and proteomes also vary dramatically across vascular beds^20, 21^. Single-cell RNA-seq has revealed new EC subpopulations defined by distinct gene expression patterns from a variety of tissues^21-23^. Although most single-cell EC profiling bins by gene expression levels, one recent report revealed the variability of VCAM1 gene expression at both RNA and protein levels in HUVEC after TNFα stimulation^24^, indicating that transcriptional heterogeneity reflects protein heterogeneity under these conditions. This heterogeneity was linked to pre-existing heterogeneity of VCAM1 promoter methylation rather than stochastic TNFα signaling or histone modification. von Willebrand factor is also expressed heterogeneously at both the RNA and protein levels, and promoter methylation was linked to control of heterogeneity in VWF expression^25^.

Here we interrogated the entire transcriptome of a cohort using three measures of variance as EC responded to homeostatic laminar flow, and we found that overall transcriptional heterogeneity increased, and was unlinked to morphological homogenization. Further analysis of selected proteins revealed that these proteins showed either no change or reduced cohort expression variance under flow, although the same RNAs increased their variance under flow, consistent with the idea that EC become more similar to each other under flow conditions. It will be interesting to determine how protein expression variance integrates with laminar flow, as both protein translation and turnover are subject to flow-mediated regulation^26, 27^. In contrast, the changes in the quantitative magnitude of RNA and protein levels of individual genes under flow relative to non-flow conditions track with each other, suggesting that the overall amount of protein, but not its variable expression within a cohort, is controlled transcriptionally by flow-mediated processes in EC. These findings have implications for EC dysfunction and disease. For example, atherosclerotic lesions form preferentially in areas of vessels that experience disturbed rather than laminar flow, such as vessel branchpoints, and EC in these areas do not exhibit the morphological homogeneity of EC in laminar areas^8, 28^. Based on our findings, we predict that important regulation of EC responses to homeostatic laminar flow includes post-transcriptional homogenization of protein levels within a cohort. Thus, therapies that work at the level of protein regulation are likely to be more effective in mitigating the effects of disturbed flow on EC function.

## ACKNOWLEDGEMENTS

We thank the UNC TGL Core for technical support for single-cell RNA-seq experiments. The authors thank members of the Bautch laboratory for constructive discussion and comments on the manuscript.

## ABBREVIATIONS

EC: endothelial cells
HUVEC: human umbilical vein endothelial cells
UMI: unique molecular identifier
RNA-seq: RNA sequencing
Gini: Gini coefficient
CV: coefficient of variation
MEF: mouse embryo fibroblasts

## HIGHLIGHTS

- Endothelial cells exposed to homeostatic laminar flow showed increased heterogeneity in RNA expression levels via single-cell profiling.
- In contrast, expression heterogeneity of cognate proteins overall decreased under laminar flow.
- Results indicate that global expression homeostasis in EC under flow is imposed post-transcriptionally.

