## supplemental Figures for "Single-Cell RNA Sequencing Reveals Endothelial Cell Transcriptome Heterogeneity under Homeostatic Laminar Flow"

Running title: EC gene expression variance under flow

[^]Current address: KBI Biopharma Inc., RTP, NC USA 27703

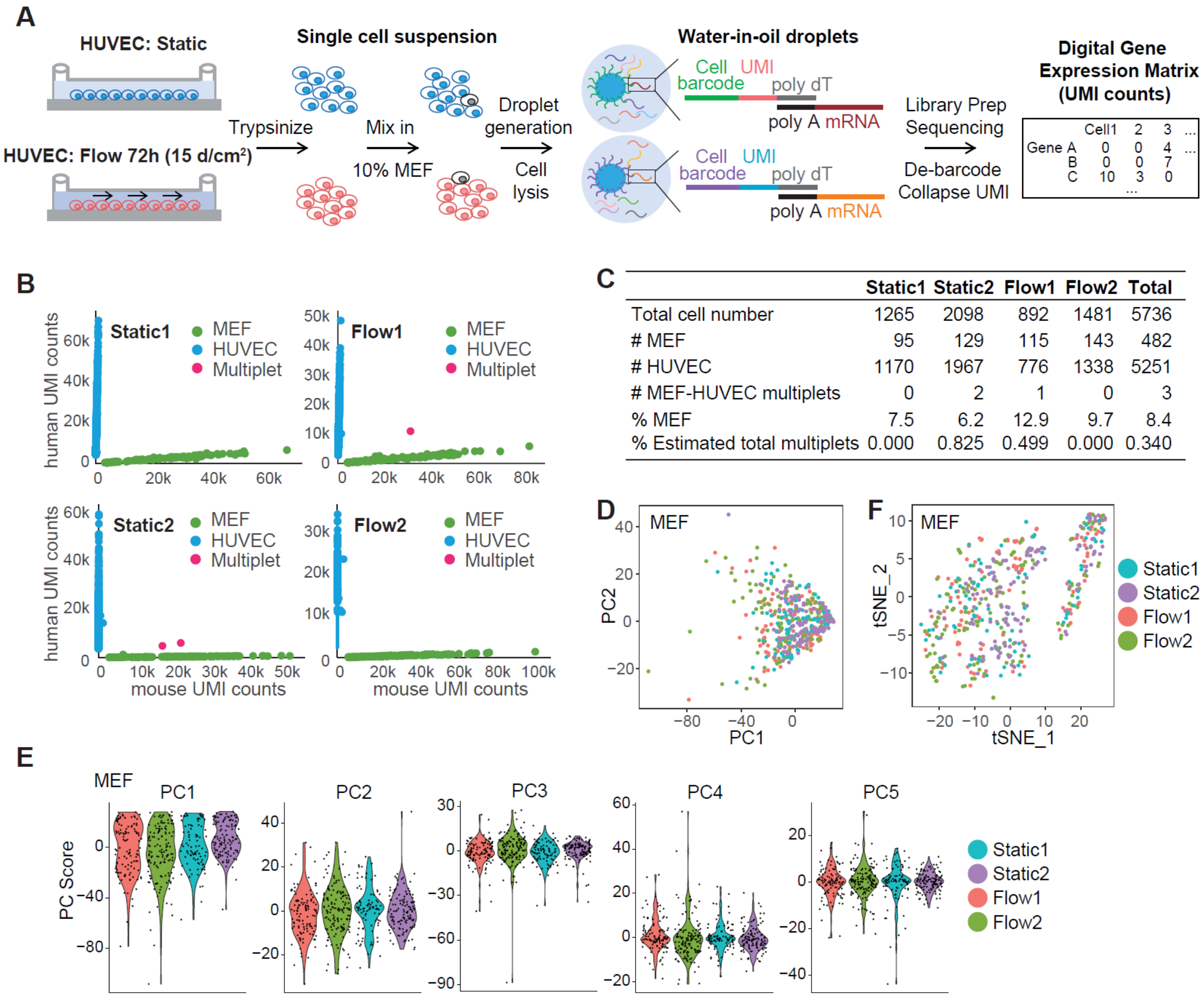


**Supplementary Figure I. Single-cell RNA-seq experiment workflow and quality control.**

(**A**) Experimental workflow. Beads are coated with numerous DNA oligos containing 14 bp cell barcode (one oligo sequence/bead, different sequences among different beads), a 10 bp unique molecular identifier (UMI, unique sequence for each oligo on the same bead), and a poly dT sequence to capture mRNA. (**B**) Multiplet discrimination of indicated samples by plotting UMI counts of each cell barcode that map to human (HUVEC, blue dots), mouse (MEF, green dots), or both (red dots) genomes. (**C**) Summary of cell number and multiplet rates. (**D**) PCA of MEF from indicated samples calculated with top 2000 highly variable genes. (**E**) Score for PC1-PC5 of PCA. (**F**) tSNE plot of MEF from indicated samples calculated with PC1-PC10.

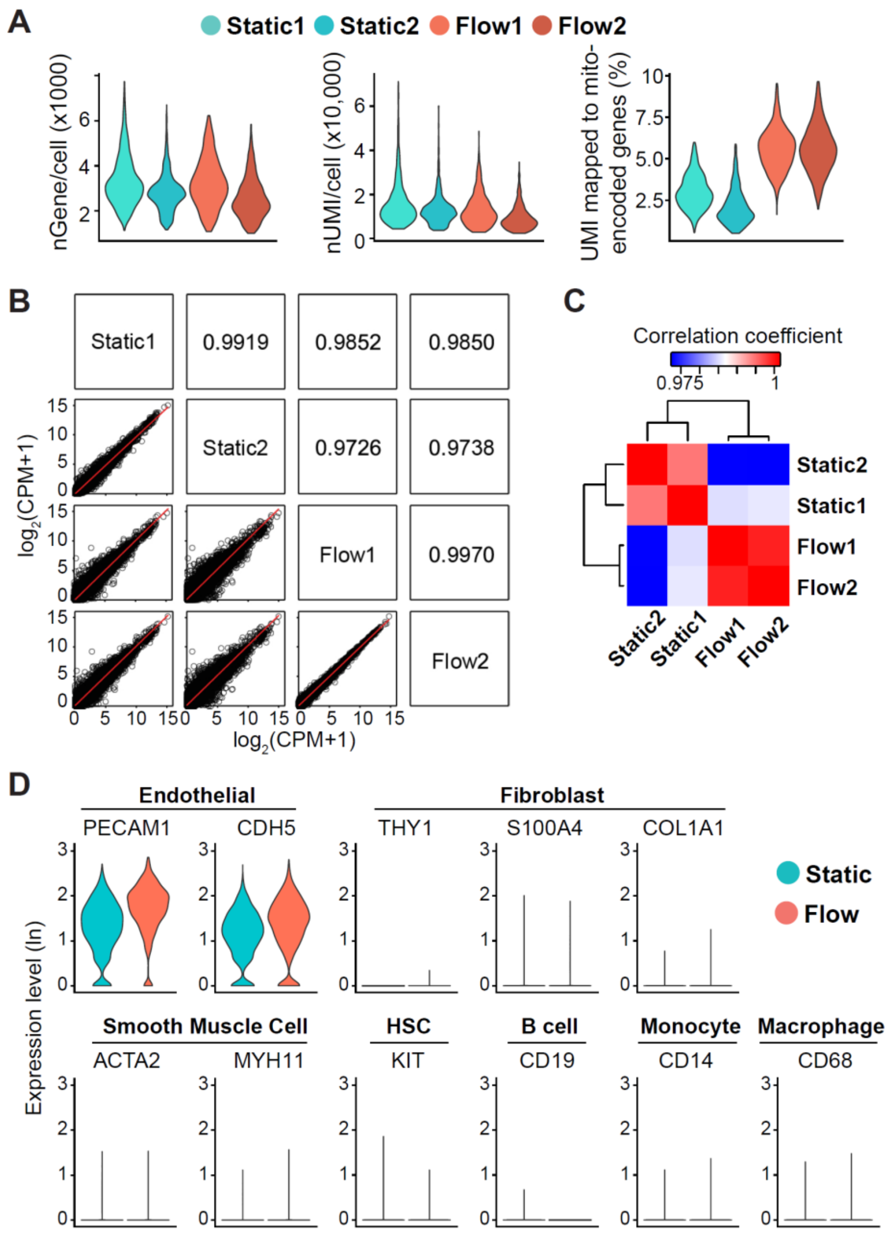


**Supplementary Figure II. HUVEC single-cell RNA-seq data quality control.**
(**A**) Distribution of detected genes/cell (left), UMI counts/cell (middle), and % UMI counts mapped to mitochondrial-encoded genes (right). (**B-C**) Pearson correlation analysis among pseudo-bulk samples assembled from the single-cell RNA-seq data showing a higher correlation between biological replicates than across conditions. (**B**) Dot plots with trend lines. (**C**) Heatmap color-coded by correlation coefficient. (**D**) Expression of canonical markers of EC, fibroblast, smooth muscle cells, hematopoietic stem cells, B cells, monocytes, and macrophages. Expression of several other markers were undetected in the dataset: CD45 (blood cells), CD235a (red blood cells), CD11b (macrophage), CD3 (T cells), CD15 (neutrophil).

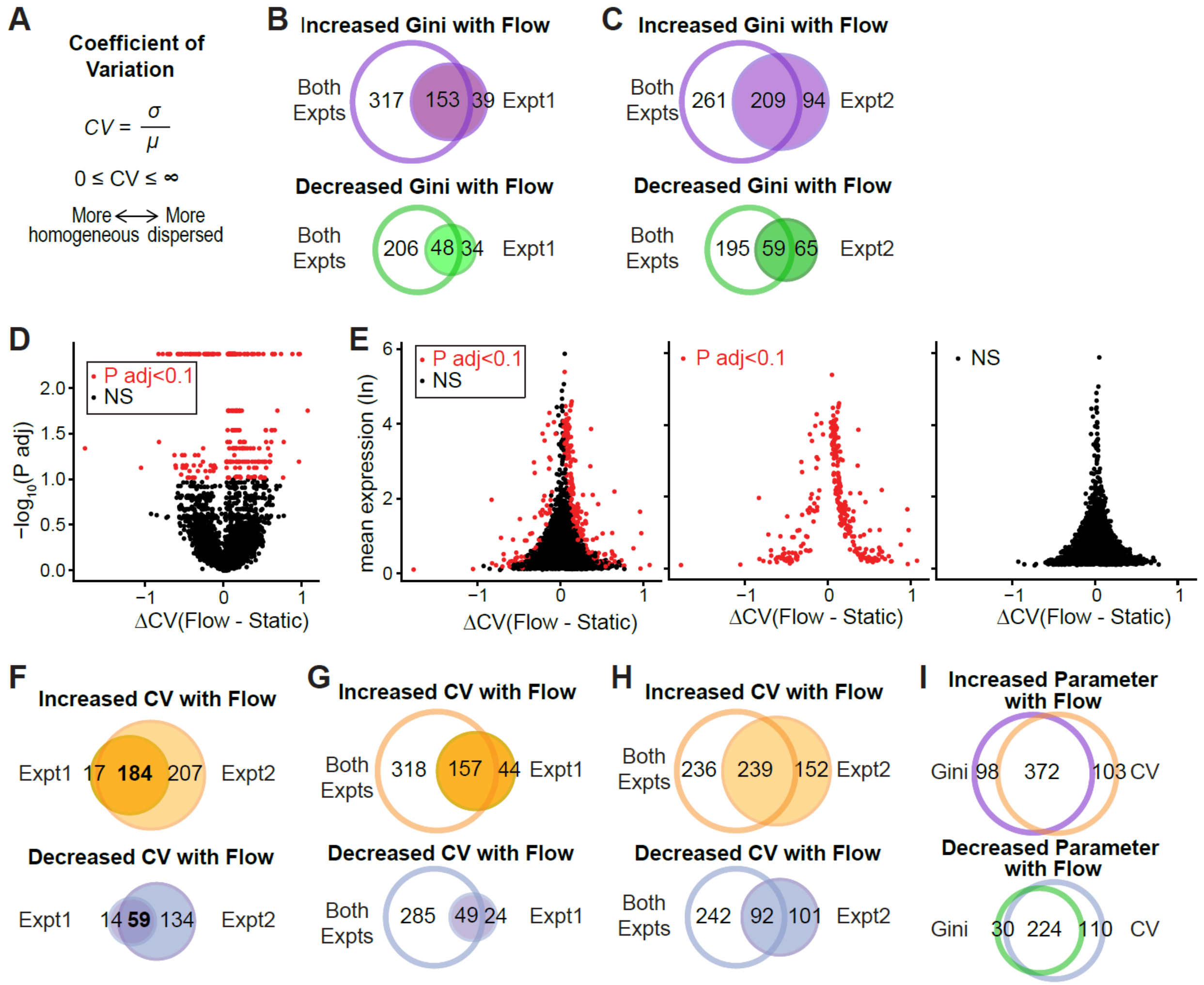


**Supplementary Figure III. Transcriptomic heterogeneity of EC under homeostatic laminar flow.**

(**A**) CV formula. (**B-C**) Differential Gini test of HUVEC transcriptomes between static and flow performed with batch as a covariate. Resulting significant gene lists were compared to results of each independent experiment without batch correction **(Fig. 1F-I)** — experiment 1 in (**B**) and experiment 2 in (**C**). (**D-G**) CV was calculated for each gene with DESCEND[13] and differential test of CV was performed for each experiment. (**D-E**) Each gene's CV change plotted against p value (**D**) or mean expression (**E**). Red dots, significant genes (P adj < 0.1), black dots, non-significant genes. (**F**) Venn plots showing overlap of genes with significant CV change in the two replicate experiments. (**G-I**) Differential CV test of HUVEC between static and flow performed for each gene with batch as a covariate. Resulting significant gene lists were compared to results of each independent experiment without batch correction — experiment 1 in (**G**) and experiment 2 in (**H**). (**I**) Venn plots showing overlap of genes with significant Gini vs. CV changes after batch correction.
